# Reduction of coastal lighting decreases seabird strandings

**DOI:** 10.1101/2023.11.15.567317

**Authors:** Tori V. Burt, Sydney M. Collins, Sherry Green, Parker B. Doiron, Sabina I. Wilhelm, William A. Montevecchi

## Abstract

Leach’s Storm-Petrels (*Hydrobates leucorhous*) are small, threatened seabirds with an extensive breeding range in the North Atlantic and North Pacific Oceans. The Atlantic population, which represents approximately 40 - 48% of the global population, is declining sharply. Positive phototaxis, the movement towards artificial light at night (ALAN), is considered to be a key contributing factor. Many seabirds exhibit positive phototaxis, which can result in stranding on land. The Leach’s Storm-Petrel is the seabird most often found stranded around ALAN in the North Atlantic, though there is little experimental evidence showing that reducing ALAN will decrease the occurrence of stranded storm-petrels. During a two-year study at a large, brightly illuminated seafood processing plant adjacent to the Leach’s Storm-Petrel’s largest colony, we compared the number of birds that stranded when the lights at the plant were turned on versus off. We recorded survival, performed carcass counts of both adults and juveniles, and released any rescued individuals. Turning the lights off reduced strandings by 39.15% (CI: 11.45% - 58.19%). The peak stranding period occurred from 25 September to 28 October, and most of the stranded birds were fledglings. These results provide evidence to support the widespread reduction and modification of coastal artificial light, especially during avian fledging and migration periods.

## Introduction

Many seabird populations have been negatively impacted by artificial light at night (ALAN) (1,2). Light from coastal towns, fishing vessels, offshore hydrocarbon platforms, and lighthouses can attract marine birds, causing them to collide with man-made structures, resulting in injury, oiling, and stranding (1,2). Stranded seabirds have difficulty taking off from land due to injury, stress, or disorientation, subjecting them to predation, starvation, and dehydration (1,3,4).

Of the seabird orders, Procellariiformes is among the most vulnerable to the effects of ALAN (2,5–8). The Leach’s Storm-Petrel (*Hydrobates leucorhous*) is one of the most nocturnally active procellariiforms (9) and is particularly vulnerable to the effects of ALAN in the North Atlantic (4,8,10). During the past 40 years, the Northwest Atlantic population of Leach’s Storm-Petrels has declined by over 50%, and as a result, the species is listed as ‘Vulnerable’ by the International Union for the Conservation of Nature (IUCN) and ‘Threatened’ by the Committee on the Status of Endangered Wildlife in Canada (COSEWIC) and the Government of Newfoundland and Labrador (11–13). The decline of the storm-petrel population is likely attributable to numerous factors, including positive phototaxis (12,14).

Little is known about why storm-petrels cluster around light, but several non-mutually exclusive and potentially interactive hypotheses have been proposed. First, birds may confuse artificial lights with bioluminescent organisms (15). One of the most important prey for Leach’s Storm-Petrels is bioluminescent myctophids (16), and it is plausible that storm-petrels are not able to differentiate between natural and artificial light (15). Second, ALAN may interfere with a storm-petrel’s ability to navigate using light from the moon and stars or their ability to sense magnetic orientation (15,17). Third, ALAN may disorient storm-petrels, causing them to be “caught” in a light catch basin from which they have difficulty escaping (1). Furthermore, light effects may be intensified in fledglings, as their retinal development is not completed until after fledging (18).

Despite the lack of understanding of why storm-petrels amass around ALAN, it has been widely suggested that turning off the lights should be used as a conservation strategy to decrease strandings (8,19). However, it is currently unknown if reducing ALAN will consequently reduce seabird strandings at common stranding sites. To our knowledge, only one study has attempted to experimentally determine the effect of ALAN on the occurrence of stranded Leach’s Storm-Petrels (6). This study, however, had very small numbers of stranded birds, and light conditions only differed between years (6), so annual differences could have accounted for the perceived effects of ALAN.

The extent to which these strandings are caused by a phototaxic response, and how that response interacts with other environmental factors, is also unclear. For example, storm-petrels may be pushed onshore by gale-force winds and become stranded (8,20). In addition, fewer storm-petrels tend to strand when moon illumination is intense (4,6,8), and on nights with greater cloud cover, more birds strand around ALAN (15). Time of year also plays a key role, as strandings tend to peak in autumn when fledglings are departing colonies and migration is ongoing (4,8,10,22).

Our research investigated intra- and inter-annual variation in the number of strandings at a major light source near the Leach’s Storm-Petrels’ largest colony on Baccalieu Island, Newfoundland and Labrador, Canada, throughout the breeding period (May - November) of 2021 and 2022. We present three hypotheses. First, we hypothesized that the stranding of storm-petrels at coastal industrial sites is a result of positive phototaxis (8,10,14,23), and we predicted that reducing ALAN by turning off the lights would reduce the number of stranded birds. Second, we hypothesized that environmental conditions influence light exposure and subsequently the amount of storm-petrel strandings around ALAN (6,8,15,20,21). We predicted that more birds would strand when wind speed and cloud cover are high, the wind is blowing onshore, and moon illumination is low. Third, we hypothesized that fledglings are more susceptible to the effects of ALAN than adults (8,10), and predicted that fledglings would be disproportionately represented in the stranded birds recovered at the site and that the peak stranding period would occur during the fledging period from early September through October (8,10).

## Methods

### Study site

Data were collected at the Quinlan Brothers Ltd. seafood processing plant in Bay de Verde, Newfoundland and Labrador, Canada (Fig 1A; 48.09780, −52.89831). Bay de Verde is within 10 km of Baccalieu Island (48.15089, −52.79700), the site of the Leach’s Storm-Petrels’ largest colony, estimated at 2 million pairs (24) (Fig 1B). Depending on weather conditions, skyward lighting from Bay de Verde can be seen from Baccalieu Island, and citizen reports indicated that thousands of storm-petrels were stranding at the plant each year, making it an ideal location to conduct this experiment. The building is 6,919 m^2^ (^25^) and has 15 bright LED lights along the top of the building facing north-west into the harbour and Conception Bay (Figs 2 and 3).

**Fig 1.**
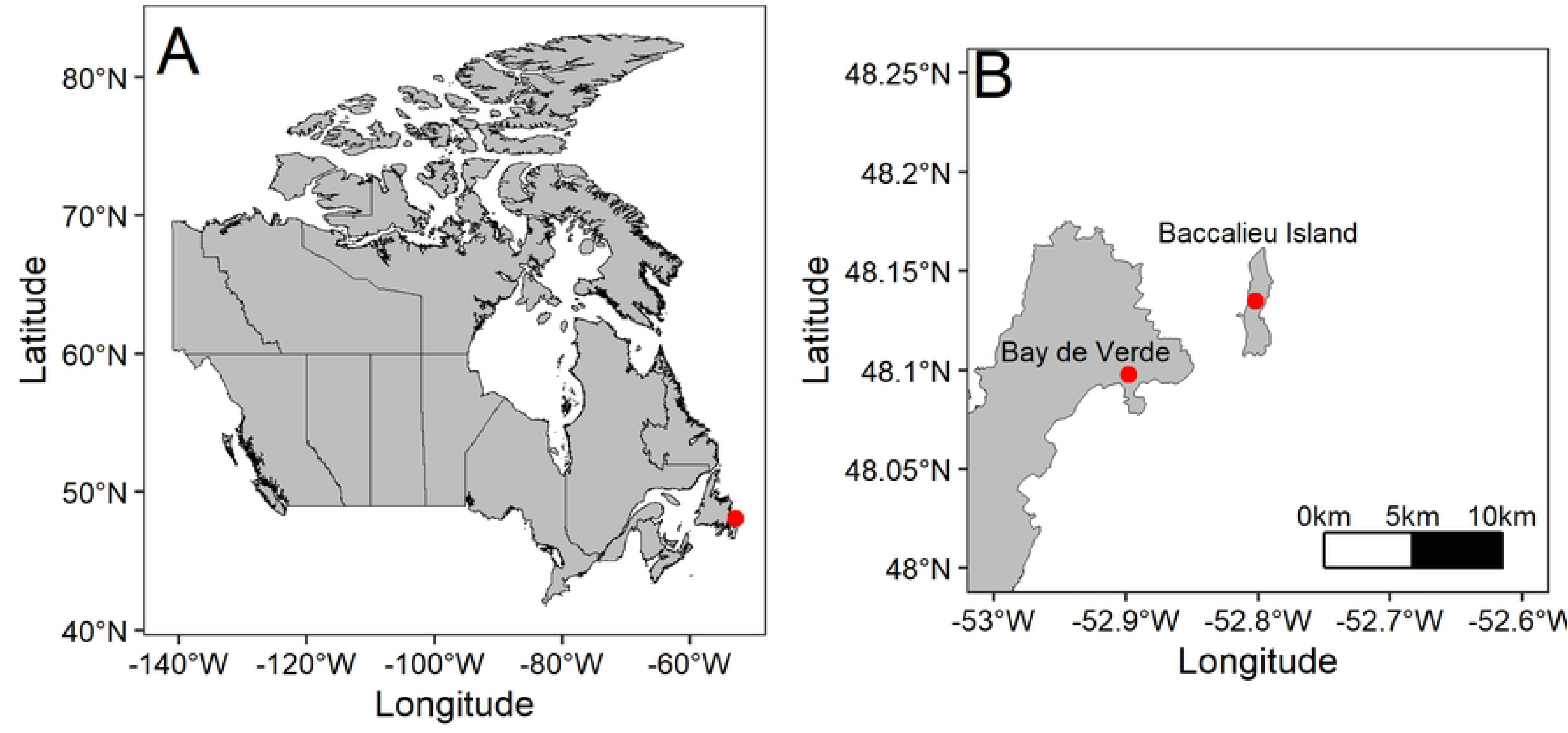
Map of Canada (A) with a red point indicating the study site Bay de Verde, Newfoundland and Labrador and (B) the study site relative to the world’s largest colony on Baccalieu Island, Newfoundland and Labrador indicated with red points.

**Fig 2.**
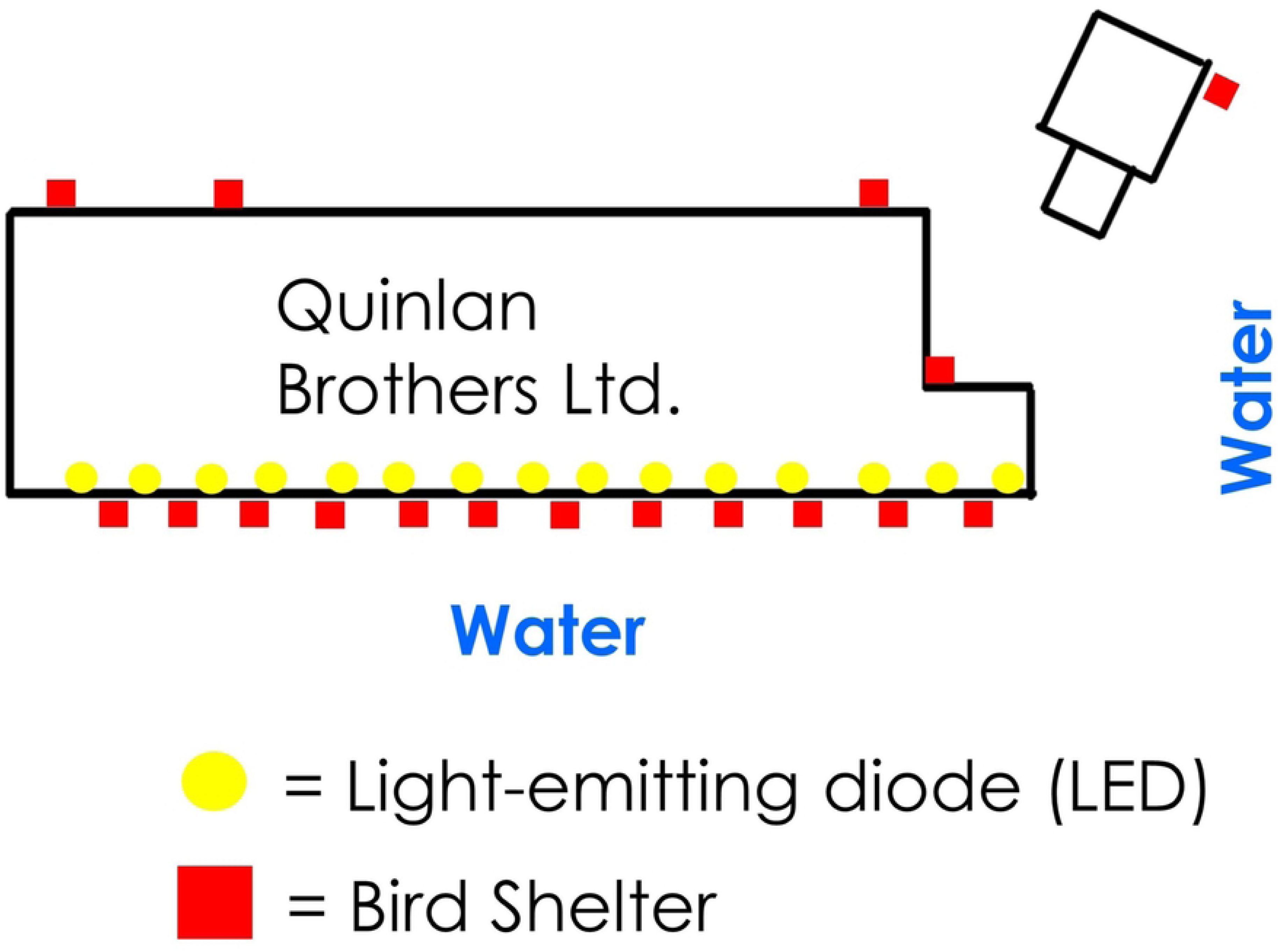
Top-down diagram of the Quinlan Brothers Ltd. seafood processing plant in 2022 (not to scale). The yellow circles indicate the LEDs on the front of the building that were being turned on and off and the red squares indicate the approximate locations of the bird shelters. The length of the front of the building is approximately 200 metres long and 10 metres high.

**Fig 3.**
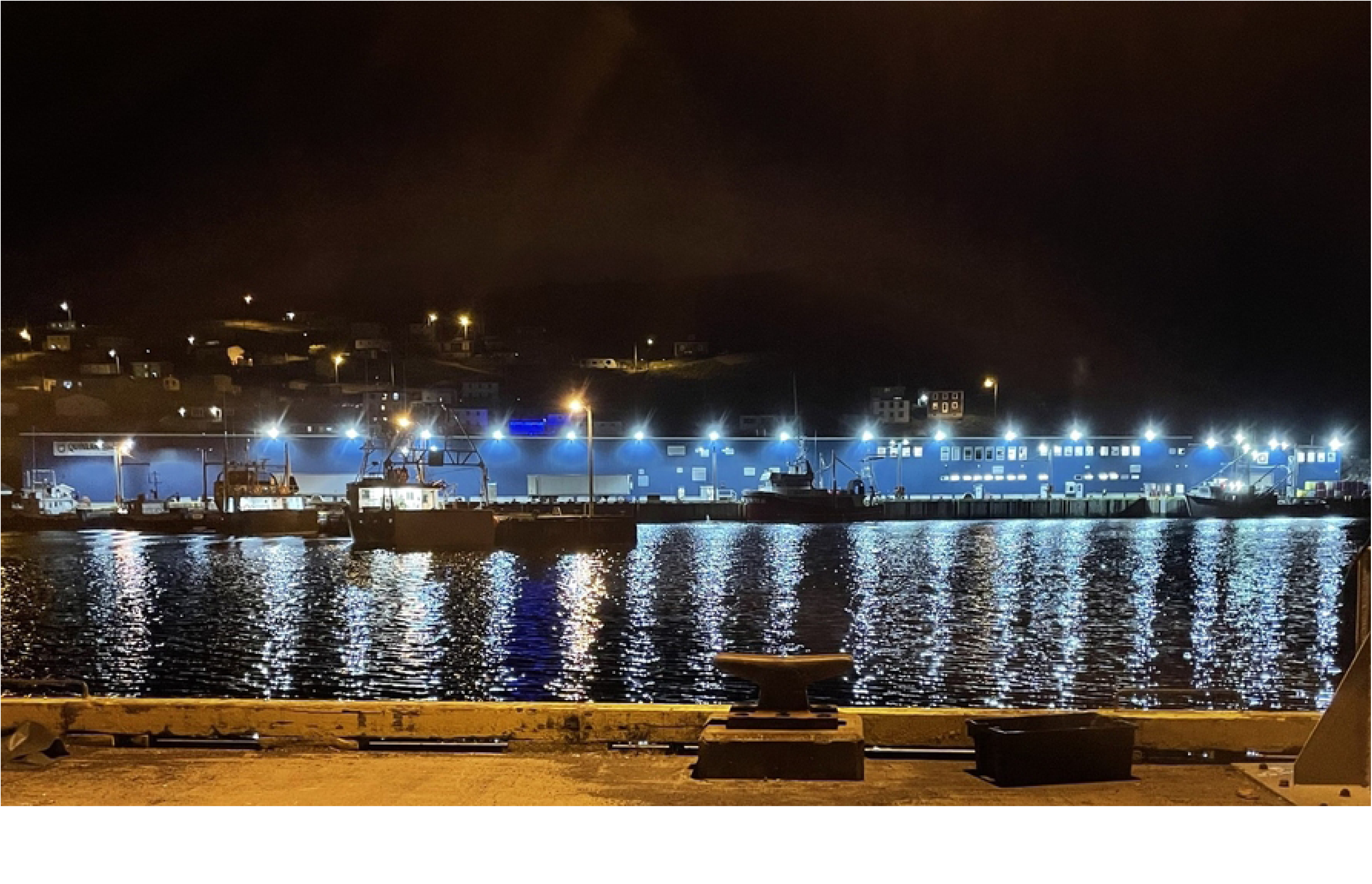
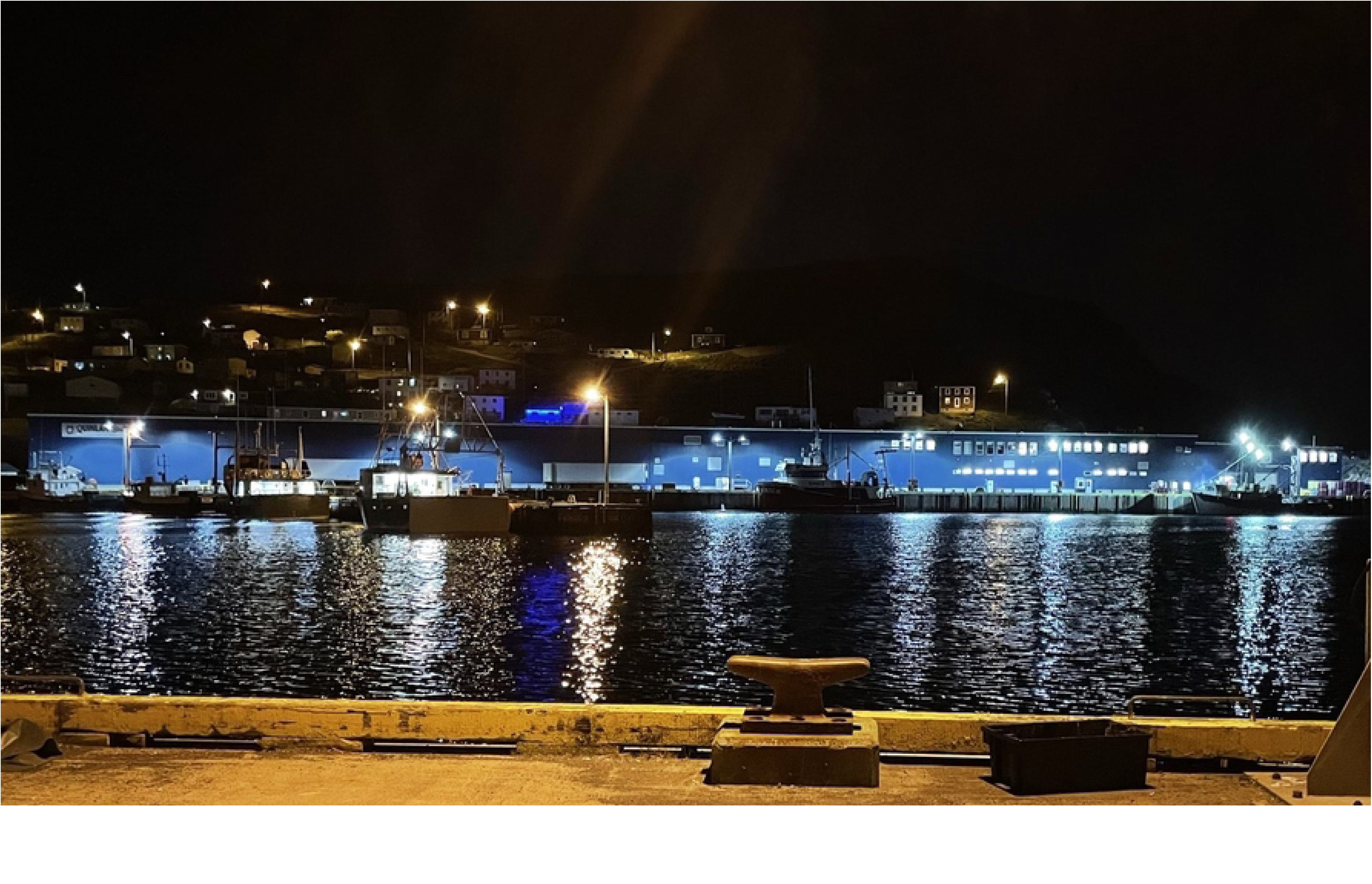
Photos taken in 2021 of the Quinlan Brothers Ltd. seafood processing plant with the LEDs on the front of the building on (A; top photo) and with nearly all of them off (B; bottom photo) [Photos taken by S.M. Collins].

### Lighting regime

During 2021, the lights on the front of the building were turned on and off according to a 28-day random schedule which aligned with the lunar cycle, for a total of 14 days of each treatment in each 28-day period. This period was chosen because reduced numbers of stranded storm-petrels have been associated with a fuller moon (8). This schedule proved difficult for the plant staff to follow, so in 2022, we implemented a schedule where the lights were turned on during odd days of the month and off during even days. This schedule was followed as closely as possible when the plant was in operation. Lights at the back of the fish plant were on at all times and lights at the front were briefly turned on, regardless of the lighting schedule, when vessels were unloading catch for safety purposes.

### Data collection

#### Daily morning surveys

Leach’s Storm-Petrels are nocturnally active burrow-nesters that seek refuge in dark confined spaces when stranded (3). In 2019, we incidentally observed Leach’s Storm-Petrels finding refuge in bait boxes used by pest control companies to control rat populations, at two industrial sites where stranding events were being investigated (this site in Bay de Verde and the Holyrood Thermal Generation Station; Wilhelm et al. 2021). As a result, we used these same boxes as bird shelters (29 x 25 x 15 cm Protecta EVO Express Bait Station after removing the inner contents) deployed around the perimeter of the building to safely capture and protect stranded birds from predation or injury. Fourteen boxes were deployed in 2021, and an additional 3 (17 total) were deployed in 2022 (Fig 2). Bird shelters were checked each morning from April 28 to November 30, 2021 and from May 3 to November 10, 2022. Surrounding areas (e.g. under concrete blocks or stairs) were also inspected and all storm-petrels found were collected. Morning checks were sometimes not completed when the plant was closed, or when nightly surveys (see below) were performed. Healthy birds were released the following evening at dusk unless poor weather conditions made it unsafe to release them. In this case, the birds were released the following evening. Injured birds were kept overnight and released the following evening if they recovered. Each morning, we recorded the date, lighting condition (on/off), weather conditions, number of live birds, number of carcasses, and number of fishing vessels that arrived at the plant. There were 158 surveys conducted in 2021 and 156 surveys conducted in 2022 (see S1 Fig. for a breakdown of surveys by month), however, some of these surveys were excluded from our analysis due to missing values (final number of daily morning surveys = 285, n_lightson_ = 111, n_lightsoff_ = 174). The lights at the plant remained off for much of September and October in 2021 and 2022, despite the lighting schedule, because the plant was not processing fish.

#### Bi-weekly night surveys

Morning checks do not capture all birds that strand due to the capacity of bird shelters, predators, and birds moving to inaccessible locations. To supplement the morning collections, nightly surveys were performed opportunistically from June 25 to October 11 2021, and 22 nightly surveys were performed approximately bi-weekly from April 20 to October 21, 2022 (S1 Fig). Starting around 22:00, we patrolled the plant perimeter every 30 - 45 minutes until around 04:00 depending on the level of bird activity. We collected live, injured, and dead storm-petrels, and checked the bird shelters and surrounding areas during each survey. Live storm-petrels were assessed for injuries, measured, weighed, aged (darker colour and condition of the primary feathers indicates juvenile birds; 26), and checked for the presence of a brood patch indicating breeding status (27). On nights when 100 - 1,000 birds were stranded (“mass stranding events”), birds were only aged and counted. Birds were released immediately from a dark area less than 1 km from the plant once assessed unless the level of stranding was high (> 50 birds) and/or the number of researchers available to rescue birds was low. When the release effort could be managed by two people, birds were accumulated in large boxes until the boxes reached capacity (~ 30 birds) and subsequently released so all researchers were available at the plant to rescue birds. If the release effort was not manageable by two people, birds were accumulated in large cardboard or fish boxes, kept in a safe place overnight, and picked up the following day to be released in Witless Bay, Newfoundland and Labrador, Canada (47.2827, −52.8323) that evening where extra help was provided to execute the release effort. Injured birds were treated in the same way as detailed above. We recorded the total number of storm-petrels found, the numbers of live and dead birds, the lighting conditions at the plant (on or off), the number of fishing vessels that arrived at the plant, and the weather conditions (cloud cover, fog, rain, and wind). There were 13 surveys conducted in 2021 and 22 surveys conducted in 2022 (see S1 Fig. for a breakdown by month), however, due to missing values for light condition, two of these had to be removed from our analysis (final number of bi-weekly night surveys = 33, n_lightson_ = 11, n_lightsoff_ = 22). The plant was shut down for approximately two to three weeks of 2022 due to extraneous circumstances, meaning the lights were off and the lighting schedule was not followed. Therefore, the lights off condition was disproportionately represented in this sample.

### Data analysis

#### Daily morning and bi-weekly night surveys

Statistical analyses and figure construction were completed using R Statistical Software version 4.2.2 (28). We assessed the period in which storm-petrels are most likely to strand by plotting the total number of stranded birds per day from the daily morning surveys in a time-series graph fit with a LOESS line of smoothing. We determined the peak stranding period as the time at which the smoothed line indicated that nightly strandings were double the mean number of stranded storm-petrels per night.

To determine whether turning the lights off at the fish plant reduced the number of stranded storm-petrels, we used a negative binomial generalized linear mixed model with a log link function. We assessed the influence of light condition (categorical; on or off), day of year, (continuous integer), year (categorical; 2021 or 2022), illuminated percent of the moon (hereafter moon illumination; continuous), wind speed (continuous), wind direction (categorical; onshore or offshore), and fog (categorical; fog or no fog) on the total number of storm-petrels stranded per night. Model construction followed the guidelines recommended by Zuur et al. (29) and Zuur and Ieno (30) and were constructed using the package “glmmTMB” (31), with assumptions tested using the package “DHARMa” (32). Moon illumination and weather information were obtained with permission from the companies Time and Date AS (33) and Custom Weather Inc (34), respectively. Wind speed and direction information were obtained from the Government of Canada website (35). Wind direction was classified as onshore or offshore based on the tangent to the eastern Newfoundland coastline from Flowers Point on the Bonavista Bay Peninsula (48.59913, −52.99618) to Sugarloaf Head on the Avalon Peninsula (47.61971, −52.65213) (S2 Fig). Mass stranding events, in which hundreds to thousands of birds strand at one site in a single evening, are relatively uncommon but can have a disproportionate effect on the model. We therefore ran the analysis both with and without the outliers (large mass stranding events of ≥ 100 birds). The inclusion of outliers did not change the inference of the model, so analyses of the whole dataset are reported.

Bi-weekly survey data were analyzed using a similar negative binomial model as the one for the daily morning surveys data, but with cloud cover (categorical, > 50% or < 50%) in place of fog, as fog data were not available for this sample. If cloud cover data were not recorded, supplementary data from Custom Weather Inc (34) were used to estimate cloud cover conditions from the closest weather station in St. Johns, Newfoundland and Labrador. During a mass stranding event in 2021, more than 1,000 birds were recorded, causing a disproportionate effect on the model. Thus, this outlier was excluded from the model.

## Results

### Effects of light and other environmental variables

#### Daily morning surveys

The number of stranded birds per night was significantly greater when the lights were turned on (df = 1, χ^2^ = 6.73, p = 0.009) and when moon illumination was low (df = 1, χ^2^ = 12.27, p = 0.0005). When the lights were off, 39.15% fewer birds tended to strand per night (CI: 11.45% - 58.19%). Predicted counts of stranded birds per night from a negative binomial generalized model were over 1.5 times higher when the lights were on compared to when the lights were off (Fig 4). Significantly more birds stranded per night in 2021 than 2022 (df = 1, χ^2^ = 10.17, p = 0.001, total birds from daily morning surveys in 2021 = 683, total birds from daily morning surveys in 2022 = 652). Day of year, wind speed and direction, and fog were not significantly associated with the number of stranded birds. Variance in the number of Leach’s Storm-Petrels stranded was significantly predicted by fog conditions (Table 1).

**Fig 4.**
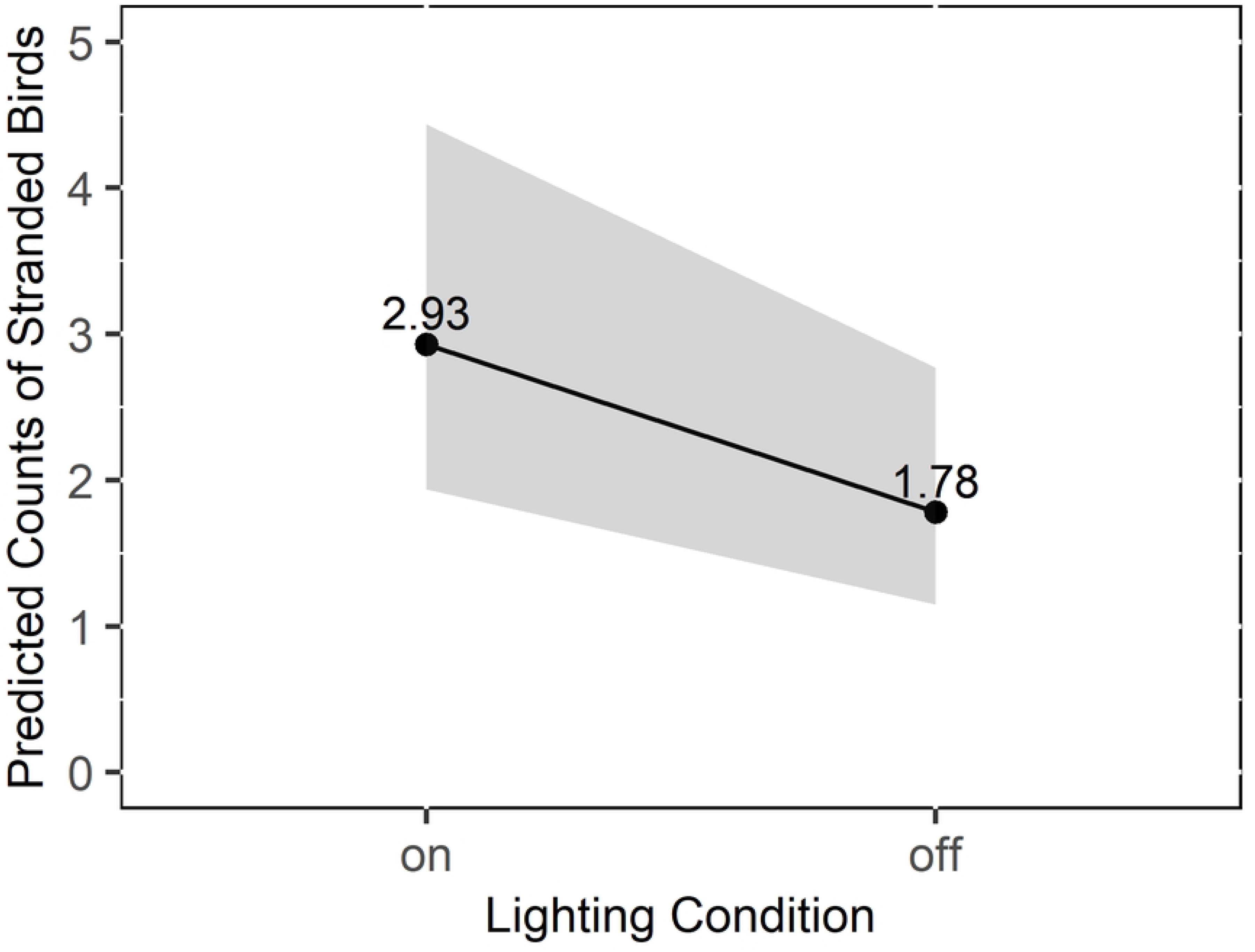
Predicted counts of Leach’s Storm-Petrels (*H. leucorhous*) strandings per night at the seafood processing plant when the lights were on versus off from a negative binomial generalized model, based on daily morning surveys. The grey outline represents the 95% confidence interval.

**Table 1.**
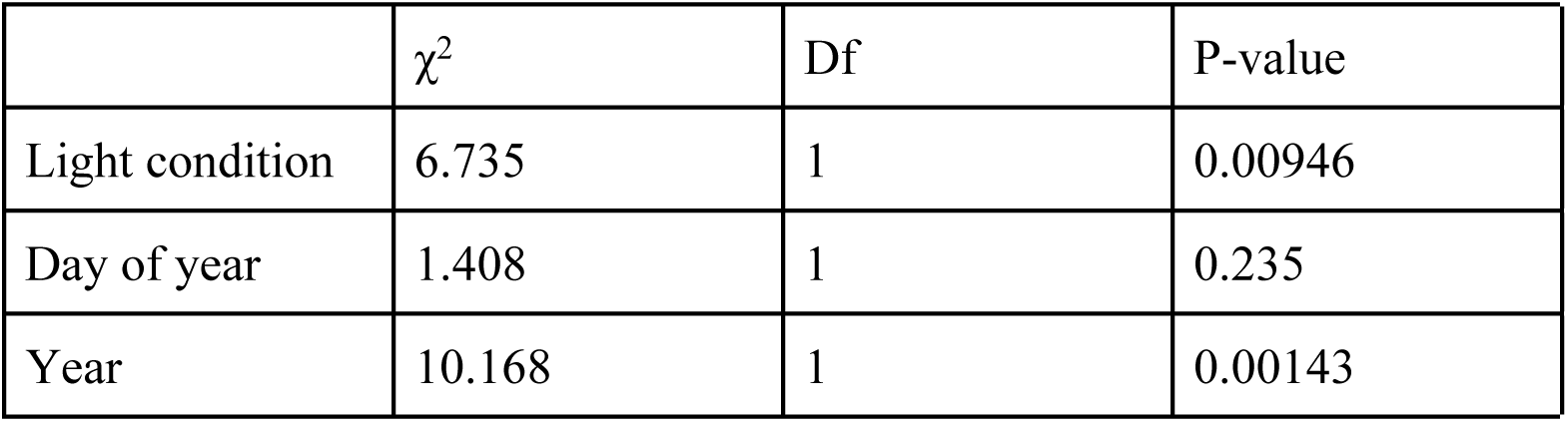
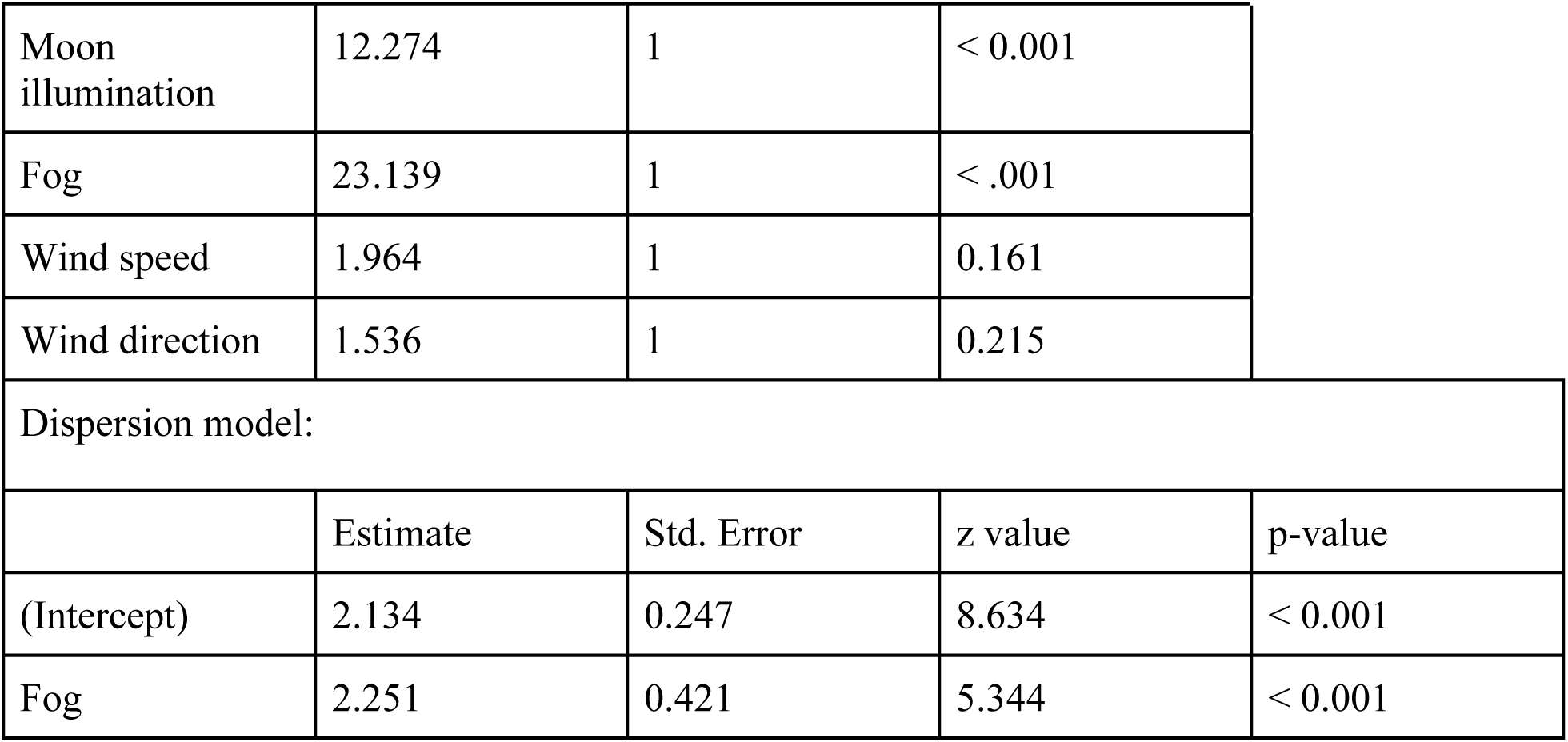
Type III ANOVA and dispersion table of the negative binomial generalized linear mixed model for the number of Leach’s Storm-Petrels (*H. leucorhous*) stranded per night at a seafood processing plant in Bay de Verde, Newfoundland and Labrador, Canada in 2021 and 2022.

| | $\chi^2$ | Df | P-value |
| --- | --- | --- | --- |
| Light condition | 6.735 | 1 | 0.00946 |
| Day of year | 1.408 | 1 | 0.235 |
| Year | 10.168 | 1 | 0.00143 |

|  |  |  |  |  |
| --- | --- | --- | --- | --- |
| Moon illumination | 12.274 | 1 | < 0.001 |  |
| Fog | 23.139 | 1 | < .001 |  |
| Wind speed | 1.964 | 1 | 0.161 |  |
| Wind direction | 1.536 | 1 | 0.215 |  |
| Dispersion model: |  |  |  |  |
|  | Estimate | Std. Error | z value | p-value |
| (Intercept) | 2.134 | 0.247 | 8.634 | < 0.001 |
| Fog | 2.251 | 0.421 | 5.344 | < 0.001 |

#### Bi-weekly night surveys

Similar to the daily morning surveys, the number of stranded birds per night during nocturnal bi-weekly surveys was significantly greater with the lights on (df = 1, χ^2^ = 7.33, p = 0.007), with less moon illumination (df = 1, χ^2^ = 3.94, p = 0.047), and during the later months of the breeding season (df = 1, χ^2^ = 14.30, p < 0.001). When the lights were off, 61.76% fewer birds tended to strand per night (CI: 23.32% - 80.93%). Predicted counts of stranded birds per night from a negative binomial generalized model were over 2.5 times higher when the lights were on compared to when the lights were off (Fig 5). Wind speed, wind direction, and cloud cover did not associate with the number of stranded storm-petrels and the number of stranded birds did not vary significantly among years. Variance in the number of Leach’s Storm-Petrels stranded was significantly predicted by the day of year (Table 2).

**Fig 5.**
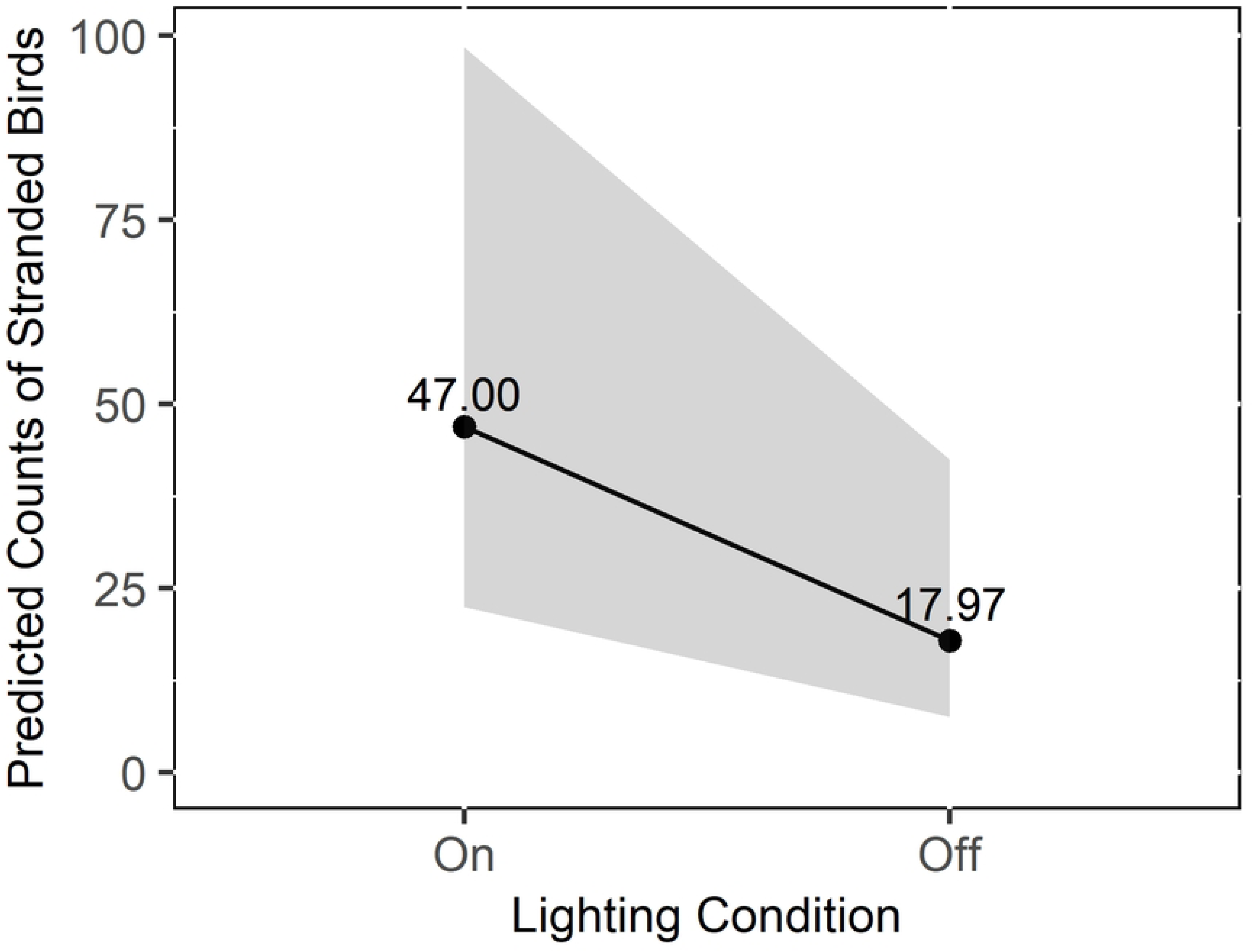
Predicted counts of Leach’s Storm-Petrels (*H. leucorhous*) strandings per night at the seafood processing plant when the lights were on versus off from the negative binomial generalized model, based on bi-weekly night surveys. The grey outline represents the 95% confidence interval.

**Table 2.** Type III ANOVA and dispersion table of the negative binomial generalized linear mixed model for the number of Leach’s Storm-Petrels (*H. leucorhous*) stranded per night at a seafood processing plant in Bay de Verde, Newfoundland and Labrador, Canada in 2021 and 2022.

|  |  |  |  |  |
| --- | --- | --- | --- | --- |
|  | Chisq | Df | p-value |  |
| Light condition | 7.331 | 1 | 0.00678 |  |
| Day of year | 14.296 | 1 | < 0.001 |  |
| Year | 0.600 | 1 | 0.438 |  |
| Moon illumination | 3.938 | 1 | 0.0472 |  |
| Cloud cover | 0.0125 | 1 | 0.911 |  |
| Wind speed | 0.879 | 1 | 0.349 |  |
| Wind direction | 0.906 | 1 | 0.341 |  |
| Dispersion model: |  |  |  |  |
|  | Estimate | Std. Error | z value | p-value |
| (Intercept) | -1.247 | 1.969 | -0.633 | 0.527 |
| Day of Year | 0.0177 | 0.00785 | 2.259 | 0.0239 |

#### Peak stranding period

Consistent with findings from the bi-weekly survey analysis that indicated a seasonal effect in the number of stranded birds, the peak stranding period occurred from September 25 to October 28 (Fig 6), when the line of smoothing indicated that the nightly strandings were consistently greater than double the mean number of birds that stranded per night (M = 4.25, SD = 27.67). This interval encompassed 81% of all birds stranded when body part estimations were included.

**Fig 6.**
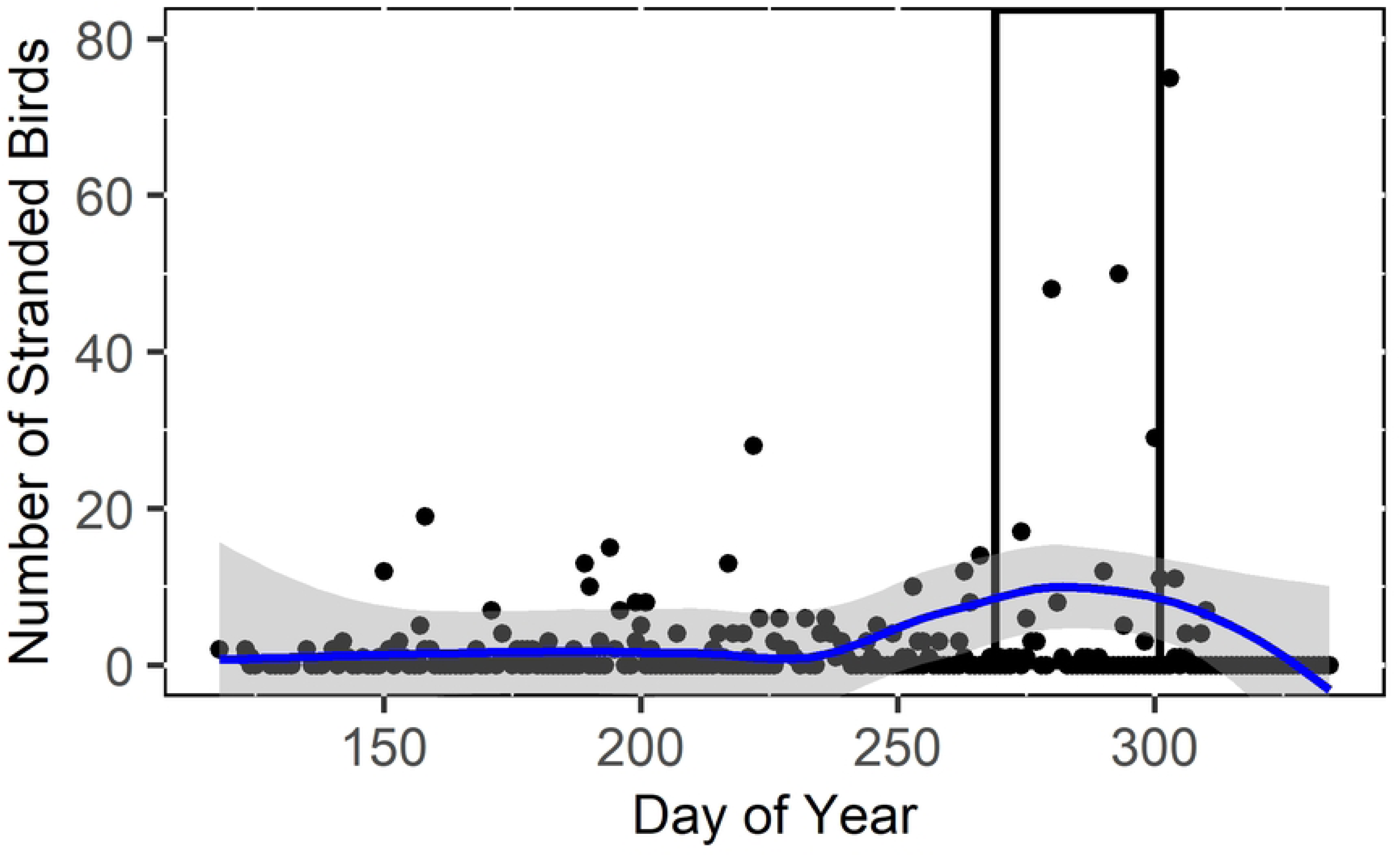
Scatterplot of the number of stranded Leach’s Storm-Petrels (*H. leucorhous*) per day of year in 2021 and 2022 collected during the daily morning surveys of a seafood processing plant in Bay de Verde, Newfoundland and Labrador, Canada (day 250 = 7 September). The blue line is the loess line of smoothing, the grey surrounding the line is the error, and the black rectangle represents the peak stranding period (September 25 - October 28). Two outliers (number of stranded birds = 387 and 283) were used to create the graph but are not shown on the above figure to improve data visualization.

### Mortality, survival, and age class

In total, including a carcass count estimation based on body part counts from September to October in 2022 (see body part count analysis in Supporting Information), 4,225 Leach’s Storm-Petrels were found stranded at the seafood processing plant in 2021 and 2022. Of these birds, 86% were alive. In 2021 and 2022, across all surveys, 1,950 birds stranded when the building LEDs were on compared to 1,929 birds when the lights were off. In total, across both years, the number of birds found during the bi-weekly night survey (n = 2,590 birds) was nearly double the number of birds found during the daily morning surveys (n = 1,335 birds). Of the birds we were able to collect and examine (live and dead), 1,609 (86.6%) were juveniles and 250 (13.4%) were adults (Table 3). Most adults stranded from June to August, and juveniles composed most (94.9%) of the birds found during the peak stranding period (Table 4).

**Table 3.**
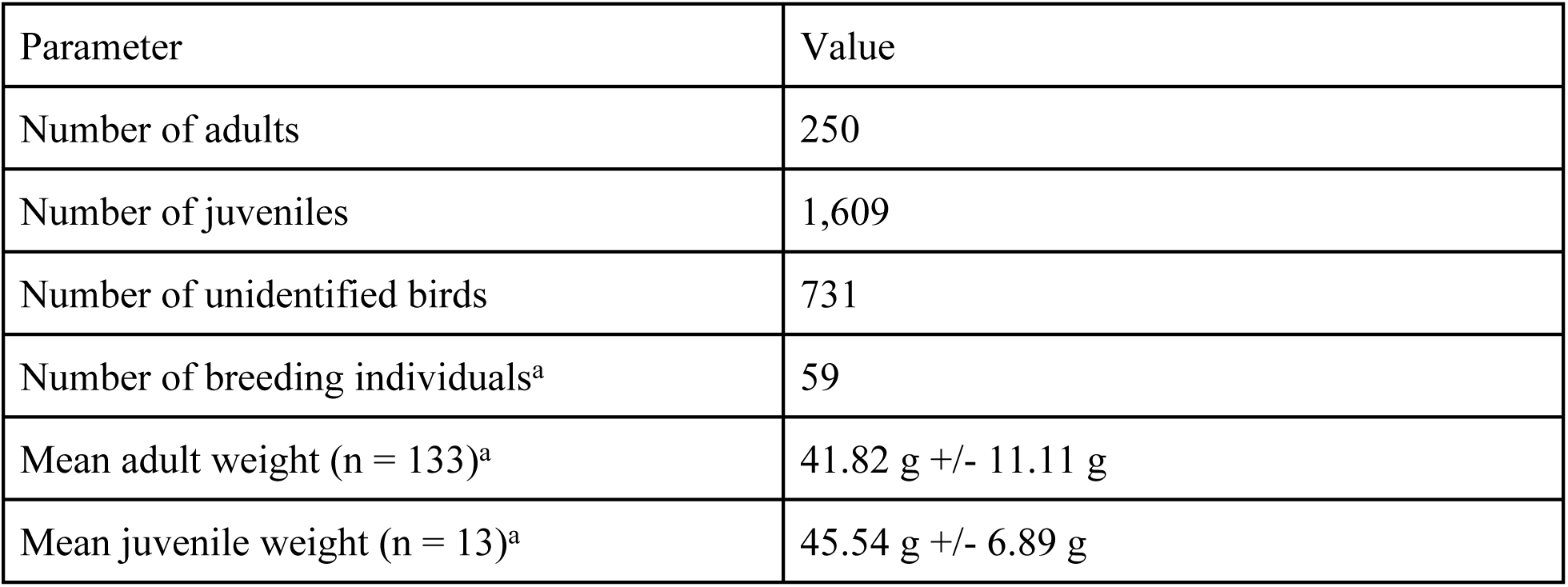
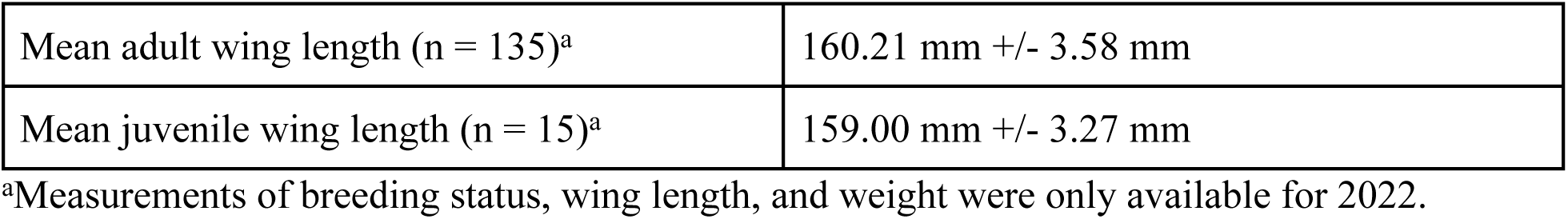
Summary of age, breeding status and individual measurements of stranded Leach’s Storm-Petrels (*H. leucorhous*) collected during bi-weekly night surveys at the seafood processing plant in Bay de Verde, Newfoundland and Labrador, Canada, during fall 2021 and throughout the 2022 breeding season.

**Table 4.** Total number of adult, juvenile, and unidentified Leach’s Storm-Petrels (*H. leucorhous*) that stranded at the seafood processing plant in Bay de Verde, Newfoundland and Labrador, Canada, per night during each month of bi-weekly survey data collection in 2021 and 2022.

|  |  | Month |  |  |  |  |  |  |
| --- | --- | --- | --- | --- | --- | --- | --- | --- |
| Year | Age | April | May | June | July | August | September | October |
| 2021 | Adult | _<sup>a</sup> | _<sup>a</sup> | 8 | _<sup>a</sup> | _<sup>a</sup> | 16 | 14 |
|  | Juvenile | _<sup>a</sup> | _<sup>a</sup> | 0 | _<sup>a</sup> | _<sup>a</sup> | 24 | 1,224 |
|  | Unidentified | _<sup>a</sup> | _<sup>a</sup> | 0 | _<sup>a</sup> | _<sup>a</sup> | 0 | 684 |
| 2022 | Adult | 0 | 0 | 46 | 41 | 70 | 9 | 46 |
|  | Juvenile | 0 | 0 | 0 | 0 | 0 | 32 | 329 |
|  | Unidentified | 0 | 0 | 0 | 0 | 0 | 8 | 39 |

<sup>a</sup>Surveys were not completed during these months.

#### Mass stranding events

Eight mass stranding events (≥ 100 birds stranding during a single night) occurred during the 2-year data collection period and all occurred within the peak stranding period; September 30 2021, October 1-4 2021, October 16, 2022, and October 19-20, 2022. Of these mass stranding events, 62.5% of mass strandings occurred when the lights were off. Notably, during 75% of these events, moon illumination only reached a high of 38%, cloud coverage was greater than 50% or it was foggy, and winds did not fall below 35 km/hr. Six of these mass stranding events occurred during the bi-weekly night survey period. During these events, juveniles comprised 66% of the birds, adults comprised only 4%, and 30% were unidentified.

## Discussion

We experimentally demonstrated that even a partial reduction in coastal lighting was an effective mitigation strategy for birds stranding around artificially lit areas, reducing strandings by 40 to 60% (Tables 1 and 2). Miles et al. (6) conducted a light experiment with Leach’s Storm-Petrels and reported that turning off the lights decreased the number of stranded birds, though their experiment demonstrated an inter-annual light effect. Our results concur with those of Miles et al. (6), but we also showed that varying the lighting condition both within and across years yields similar findings, suggesting that Leach’s Storm-Petrels exhibit positive phototaxis. Our results can likely be extended to related species. It is well established that other burrow-nesting procellariiforms cluster around ALAN (5,7,36,37). For example, Telfer et al. (15) found more than 1,000 grounded Newell’s Shearwaters (*Puffinus newelli*) each year during an experiment on Kauai Island, Hawaii from 1978 to 1985. In 1982 and 1983, 79% of those stranded shearwaters were recovered in the brightly illuminated urban areas of the island (15). On Phillip Island in Australia, more than 8,000 Short-tailed Shearwaters (*Ardenna tenuirostris*) were grounded from 1999 to 2013, suggesting that they also exhibit positive phototaxis (7). This population of shearwaters also responded positively to reduced ALAN; turning off the lights on a brightly illuminated bridge decreased the number of shearwaters stranded on the bridge (7).

In addition to ALAN, moon illumination was a key environmental factor which influenced strandings. Fewer storm-petrels stranded when moon illumination was high (Tables 1 and 2). Cloud cover data were not available so inferences about moon brightness in the sky, as perceived by the storm-petrels, cannot be made. Numerous studies show that greater moon illumination is associated with decreased strandings and reduced colony activity by Leach’s Storm-Petrels and other procellariiforms (4,5,8,15,19). This phenomenon has two proposed explanations: 1) lunar illumination appears to reduce the disorienting influence of ALAN (19), and 2) birds are less active on nights with high moon illumination (38), and as a result, fewer strandings occur. A recent study by Collins et al. (39) supports the first hypothesis. Although adults tend to be less active on a full moon (38), Leach’s Storm-Petrel chick fledging was not associated with moon phase or incident illumination (39). Given that fewer birds in our study stranded on a full moon and that most collected in our study were fledglings (Table 3), it is likely that moonlight reduces the risk for stranding by reducing the attractive properties of ALAN (21,39). Notably, almost all mass stranding events occurred when moon illumination was low, and cloud cover was high or it was foggy. Though fog, cloud cover, and wind speed and direction did not associate with the number of stranded birds, contradicting previous research (20), these results suggest that fog and cloud cover, environmental variables which influence available nocturnal illumination, may play a role in mass stranding events. Interestingly, most mass stranding events (63%) occurred when the lights were off. This proportion is similar to the proportion of nights in our study when the lights were off (61%), suggesting that mass stranding events occur randomly relative to the light condition at the plant. To make better-informed assessments of the effect of cloud cover and fog, more research is needed into these episodic occurrences.

In the 2021 and 2022 breeding seasons, researchers and volunteers rescued over 3,500 stranded Leach’s Storm-Petrels at the seafood processing plant in Bay de Verde. Despite daily monitoring, hundreds of birds perished around the plant. Our results demonstrate that the number of birds found when completing nightly surveys was almost double the number found when conducting morning surveys, even though the bi-weekly night surveys only accounted for 10% of the total surveying effort (S1 Fig). Daily morning surveys, while often the only feasible option, do not account for birds that were removed by predators or died in inaccessible locations. To maximize the survival of stranded Leach’s Storm-Petrels, consistent researcher or volunteer presence is needed at common stranding areas during the peak stranding period to conduct nightly surveys. The peak stranding period of birds at Bay de Verde occurred from September 25 to October 28 (Fig 6) and bi-weekly surveys showed that the number of stranded birds increased through the season (Table 2). These results concur with previous research which found that most storm-petrels stranded during late September through to November (3,4,6,8,10). This period corresponds directly with the Leach’s Storm-Petrels’ fledging period (27,39) and most birds found during this period were fledglings (Table 4), suggesting that they are the age class most susceptible to the effects of ALAN (15,18,40). These stranding events consisted of mostly fledglings but seemed to occur randomly relative to light conditions at the plant (62.5% occurred when the building lights were off). These results can be interpreted as support for one of two hypotheses: 1) fledglings exhibit stronger positive phototaxis relative to adults, and a partial reduction in lighting is not enough to reduce strandings, or 2) fledglings exhibit weaker positive phototaxis relative to adults. Our results, along with previous findings, concur with the first hypothesis, as birds stranded near sources of ALAN in September and October, many of which were identified as or presumed to be fledglings (8,10). It should be noted that the proportion of mass stranding events that occurred with the lights off is similar to the proportion of lights off nights included in our sample (196 total number of lights off nights during the daily morning and bi-weekly night surveys/318 total number of daily morning and bi-weekly night surveys, 61.6%).

While the loss of fledglings is concerning, the hundreds of adults, 59 of which were verified breeders based on brood patch presence (Table 3), raises concerns for population-level effects. Most adults that were verified as breeders stranded from June to August. The number of stranded breeding birds may have been underestimated, as the brood patch feathers begin to re-grow four weeks after hatching (27), making it difficult to determine breeding status after this period. The mortality of breeding adults is concerning because storm-petrels exhibit delayed maturity, breed once a year, and lay a single egg annually (27). When a breeding adult dies, it disrupts a long-term monogamous pair bond (27) and may impact the health and survival of the egg or chick, as seen in other long-lived monogamous seabird species (41). Though individuals can form new pair bonds, new pairs may take time to re-establish and often have poor reproductive success for several years (42). Even though most birds strand from September to October, it is important to reduce nocturnal lighting throughout the breeding season to protect breeding adults as well as juveniles.

Knowledge of the timeline and locations of when and where storm-petrels are most likely to strand allows for a fine-tuning of conservation efforts related to positive phototaxis. A recent study examined social media reports of Leach’s Storm-Petrel strandings across the island of Newfoundland and found that the majority of birds stranded in brilliantly illuminated urban areas (10), suggesting that ALAN is likely causing widespread strandings throughout insular Newfoundland and the species’ breeding range [see also (8)]. Conservation efforts at the seafood processing plant, including turning off the lights, using bird shelters, and searching daily for stranded birds to reduce mortality are effective elsewhere (e.g., Newfoundland and Labrador Hydro Thermal Generation Station), though these efforts are not always feasible. For example, vessels arrive throughout the night at the plant in Bay de Verde and the lights must be on to ensure safe visibility when unloading catch. Furthermore, bird shelters must be checked daily to retrieve live birds. Therefore, it is important to seek out other permanent solutions to reduce strandings by exploring the properties of light that influence phototaxis such as wavelength, light source, intensity, and the size of the light catch basin (5,43,44). Contention surrounds the attractive properties of different wavelengths of light. Some research suggests that blue and green light attract the most migratory birds in cloudy conditions (45), while other results suggest that red and white light attract the most seabirds and migratory birds and cause the most disorientation in foggy conditions (43,46). The type of light source can also have an impact; high-pressure sodium lights attract fewer shearwaters than metal halide lights and LEDs (44). Specific research investigating the wavelengths and light types to which Leach’s Storm-Petrels are the most attracted has not been conducted. Future research should focus on determining the spectral sensitivities of adult LESP [but see Mitkus et al. (40)] to inform decisions about mitigations to reduce population mortality. Shielding light has been successful for limiting attraction to ALAN by other procellariiforms, because shielding redirects the light to reduce the light catch basin projected skyward (5). Similarly, motion-activated lights should be considered in areas where constant lighting is unnecessary.

Increasing rescue efforts at hot-spot stranding locations where near zero-lighting cannot be achieved is also recommended to maximize recovery of stranded storm-petrels. Performing systematic nightly searches during the peak stranding period (from late September to the end of October) and deploying an array of bird shelters at the start of the breeding season (early May) that are checked daily can reduce mortality of breeding adults and recently fledged juveniles at common stranding locations. This strategy is currently employed by the “Puffin and Petrel Patrol” (47), a non-profit organization that provides volunteers with training on how to rescue stranded fledgling puffins and storm-petrels, and subsequently sends volunteers to common stranding locations in southeastern Newfoundland during the night. Storm-petrels strand across a large geographical scale (4,8,10) and widespread public education regarding light-induced storm-petrel strandings coupled with the development of volunteer rescue groups throughout their breeding range could dramatically reduce seabird mortality associated with attraction to ALAN.

## Conclusion

The evidence indicates that turning off extraneous lights can significantly decrease Leach’s Storm-Petrel strandings and that even partial light reductions are effective in this regard. More than 4,000 storm-petrels were found stranded at a seafood processing plant in Newfoundland and Labrador in 2021 and 2022. This stranding estimate is likely highly conservative, as it does not include mortality that occurred throughout the beginning of the breeding season when researchers were not present, nor does it include mortality that occurred throughout the night as a result of predators removing evidence of predation on birds that could not find a safe place to hide. Increasing research and rescue efforts and promoting public education about Leach’s Storm-Petrels is important, but employing a preventative strategy to mitigate seabird strandings is a more efficient and effective solution. Therefore, we recommend shifting conservation efforts to focus on reducing unnecessary extraneous sources of ALAN as much as possible, particularly during late September through October.

## Acknowledgments

We thank Robin Quinlan, Barry Hatch, Kristinn Skulason, Ed and Cindy Noonan, Jim and Cheryl Broderick, and all of the staff at the Quinlan Brothers Ltd. seafood processing plant for permitting and supporting the project, and for assisting with the light schedule and bird collection. Thank you to Robert Blackmore, Taylor Brown, Juliana Coffey, Kyle D’Entremont, Mohammad Fahmy, Gretchen McPhail, Fiona Le Taro, and Christopher Ward for help collecting and releasing birds. Thank you to Alex Day for assistance with data analysis. We thank Time and Date AS and Custom Weather Inc. for providing weather data. We are grateful to Jake Russell-Mercier of ECCC for continued diligence and efforts to help fund this research.

## Ethical considerations

Birds were collected under Canadian Wildlife Service scientific permit no. LS2688, and banded under Environment and Climate Change Canada Scientific Permits to Capture and Band Migratory Birds no. 10559 X and 10332 K.

## Notes

### Competing Interest Statement

The authors have declared no competing interest.

